# The Ionotropic Receptors IR21a and IR25a mediate cool sensing in *Drosophila*

**DOI:** 10.1101/032540

**Authors:** Lina Ni, Mason Klein, Kathryn Svec, Gonzalo Budelli, Elaine C. Chang, Richard Benton, Aravinthan D. T. Samuel, Paul A. Garrity

## Abstract

Animals rely on highly sensitive thermoreceptors to seek out optimal temperatures, but the molecular mechanisms of thermosensing are not well understood. The Dorsal Organ Cool Cells (DOCCs) of the *Drosophila* larva are a set of exceptionally thermosensitive neurons critical for larval cool avoidance. Here we show that DOCC cool-sensing is mediated by Ionotropic Receptors (IRs), a family of sensory receptors widely studied in invertebrate chemical sensing. We find that two IRs, IR21a and IR25a, are required to mediate DOCC responses to cooling and are required for cool avoidance behavior. Furthermore, we find that ectopic expression of IR21a can confer cool-responsiveness in an *Ir25a-*dependent manner, suggesting an instructive role for IR21a in thermosensing. Together, these data show that IR family receptors can function together to mediate thermosensation of exquisite sensitivity.

## INTRODUCTION

Temperature is an omnipresent physical variable with a dramatic impact on all aspects of biochemistry and physiology(Sengupta and Garrity, 2013). To cope with the unavoidable spatial and temporal variations in temperature they encounter, animals rely on thermosensory systems of exceptional sensitivity (Barbagallo and Garrity, 2015; Dhaka et al., 2006). These systems are used to avoid harmful thermal extremes and to seek out and maintain body temperatures optimal for performance, survival and reproduction (Barbagallo and Garrity, 2015; Flouris, 2011).

Among the most sensitive biological thermoreceptors described to date are the Dorsal Organ Cool Cells (DOCCs), a recently discovered trio of cool-responsive neurons found in each of the two dorsal organs at the anterior of the *Drosophila melanogaster* larva (Klein et al., 2015). The DOCCs robustly respond to decreases in temperature as small as a few milli-degrees C per second (Klein et al., 2015), a thermosensitivity similar to that of the rattlesnake pit organ (Goris, 2011), a structure known for its extraordinary sensitivity. At the behavioral level, the DOCCs are critical for mediating larval avoidance of temperatures below ~20°C, with the thermosensitivity of this avoidance behavior paralleling the thermosensitivity of DOCC physiology (Klein et al., 2015). While the DOCCs are exceptionally responsive to temperature, the molecular mechanisms that underlie their thermosensitivity are unknown.

At the molecular level, several classes of transmembrane receptors have been shown to participate in thermosensation in animals. The most extensively studied are the highly thermosensitive members of the Transient Receptor Potential (TRP) family of cation channels (Clapham and Miller, 2011; Dhaka et al., 2006). These TRPs function as temperature-activated ion channels and mediate many aspects of thermosensing from fruit flies to humans (Barbagallo and Garrity, 2015; Damann et al., 2008; Dhaka et al., 2006). In addition to TRPs, other classes of channels contribute to thermosensation in vertebrates, including the thermosensitive calcium-activated chloride channel Anoctamin 1 (Cho et al., 2012) and the two pore potassium channel TREK-1 (Alloui et al., 2006). Recent work in *Drosophila* has demonstrated that sensory receptors normally associated with other modalities, such as chemical sensing, can also make important contributions to thermotransduction. In particular, GR28B(D), a member of the invertebrate gustatory receptor (GR) family, was shown to function as a warmth receptor to mediate warmth avoidance in adult flies exposed to a steep thermal gradient (Ni et al., 2013). The photoreceptor Rhodopsin has also been reported to contribute to temperature responses, although its role in thermosensory neurons is unexamined (Shen et al., 2011).

Ionotropic Receptors (IRs) are a family of invertebrate receptors that have been widely studied in insect chemosensation, frequently serving as receptors for diverse acids and amines (Silbering et al., 2011). The IRs belong to the ionotropic glutamate receptor (iGluR) family, an evolutionarily conserved family of heterotetrameric cation channels that includes critical regulators of synaptic transmission like the NMDA and AMPA receptors (Croset et al., 2010). In contrast to iGluRs, IRs have only been found in Protostomia and are implicated in sensory transduction rather than synaptic transmission (Rytz et al., 2013). In insects, the IR family has undergone significant expansion and diversification, with the fruit fly *D. melanogaster* genome encoding 66 IRs (Croset et al., 2010). While the detailed structures of IR complexes are unknown, IRs are often thought to form heteromeric channels in which an IR “co-receptor” (such as IR25a, IR8a or IR76b) partners one or more “stimulus-specific” IRs (Abuin et al., 2011).

Among insect IRs, IR25a is the most highly conserved across species (Croset et al., 2010). In *Drosophila*, IR25a expression has been observed in multiple classes of chemosensory neurons with diverse chemical specificities, and IR25a has been shown to function as a “co-receptor” that forms chemoreceptors of diverse specificities in combination with other, stimulus-specific IRs (Abuin et al., 2011; Rytz et al., 2013). IR21a is conserved in mosquitoes and other insects, but has not been associated with a specific chemoreceptor function (Silbering et al., 2011), raising the possibility that it may contribute to other sensory modalities.

Here we show that the previously “orphan” IR, *Ir21a*, acts together with the co-receptor IR25a to mediate thermotransduction. We show that these receptors are required for larval cool avoidance behavior as well as the physiological responsiveness of the DOCC thermosensory neurons to cooling. Furthermore, we find that ectopic expression of IR21a can confer cool responsiveness in an *Ir25a*-dependent manner, indicating that IR21a can influence thermotransduction in an instructive fashion.

## RESULTS

### Dorsal organ cool cells express *Ir21a-Gal4*

To identify potential regulators of DOCC thermosensitivity, we sought sensory receptors specifically expressed in the dorsal organ housing these thermoreceptors (Fig. 1a). Examining a range of potential sensory receptors in the larva, we found that regulatory sequences from the Ionotropic Receptor Ir21a drove robust gene expression (via the Gal4/UAS system (Brand and Perrimon, 1993)) in a subset of neurons in the dorsal organ ganglion (Fig. 1b, 1c). We observed *Ir21a-Gal4* drove gene expression in three neurons within each dorsal organ ganglion (Fig. 1b, 1c). These neurons exhibited the characteristic morphology of the DOCCs, which have unusual sensory processes that form a characteristic “dendritic bulb” inside the larva (Klein et al., 2015).

**Figure 1:**
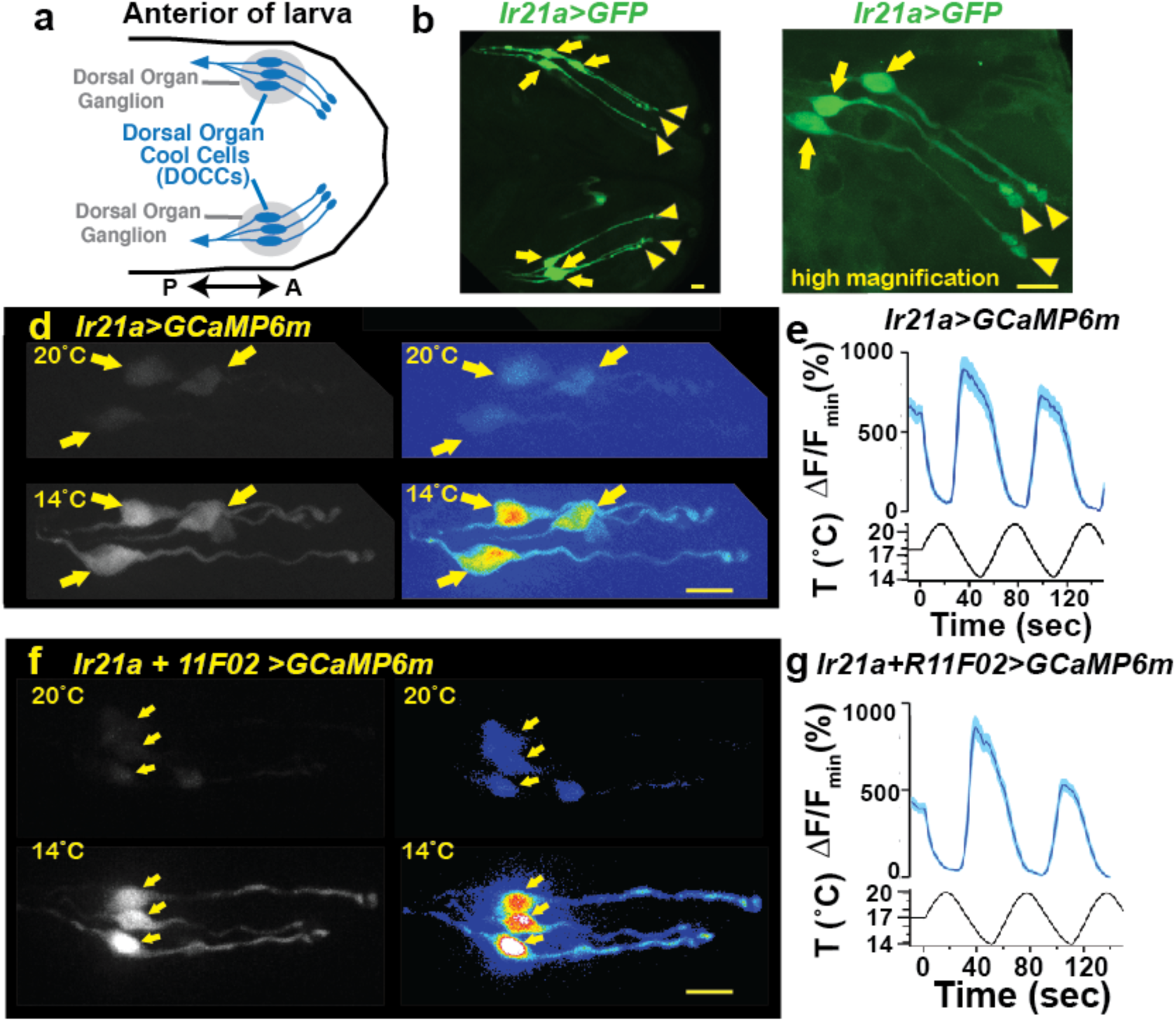
Dorsal Organ Cool Cells (DOCCs) express *Ir21a-Gal4*. a) First/second instar larval anterior. Each Dorsal Organ Ganglion (grey) contains three DOCCs (blue). Anterior-Posterior axis denoted by double-headed arrow. b,c) *Ir21a-Gal4;UAS-GFP* (*Ir21a>GFP*) labels larval DOCCs. Arrows denote cell bodies and arrowheads dendritic bulbs. d) Temperature responses of *Ir21a-Gal4;UAS-GCaMP6m*-labeled DOCCs. Left panels, raw images; right panels, colors reflect fluorescence intensity. Arrows denote cell bodies. e) Fluorescence quantified as percent change in fluorescence intensity compared to minimum intensity. n=22 cells. f,g) Temperature-responses of *Ir21a-Gal4;R11F02-Gal4;UAS-GCaMP6m*-labeled DOCCs. n=26. Scale bars, 10 microns. Traces, average +/− SEM.

To confirm that the *Ir21a*-*Gal4*-positive neurons were indeed cool-responsive, their thermosensitivity was tested by cell-specific expression of the genetically encoded calcium indicator GCaMP6m under *Ir21a-Gal4* control. Consistent with previously characterized DOCC responses (Klein et al., 2015), when exposed to a sinusoidal temperature stimulus between ~14°C and ~20°C, GCaMP6m fluorescence in these neurons increased upon cooling and decreased upon warming (Fig. 1d, 1e and Supp. Fig. 1). The expression of *Ir21a-Gal4* was also compared with that of *R11Fo2-Gal4*, a promoter used in the initial characterization of the DOCCs (Klein et al., 2015). As expected, GCaMP6m expressed under the combined control of *Ir21a-Gal4* and *R11Fo2-Gal4* revealed their precise overlap in three cool-responsive neurons with DOCC morphology in the dorsal organ, further confirming the identification of the *Ir21a-Gal4-*expressing cells as the cool-responsive DOCCs (Fig. 1f,g).

### *Ir21a* mediates larval thermotaxis

To assess the potential importance of *Ir21a* in larval thermosensation, we tested the ability of animals to thermotax when *Ir21a* function has been eliminated. Two *Ir21a* alleles were generated, *Ir21a^123^* and *Ir21a*^Δ^*^1^. Ir21a^123^* deletes 23 nucleotides in the region encoding the first transmembrane domain of IR21a and creates a translational frameshift (Fig. 2a). *Ir21a*^Δ^*^1^* is an ~11 kb deletion removing all except the last 192 nucleotides of the *Ir21a* open reading frame, including all transmembrane and ion pore sequences (Fig. 2a). As the deletion in *Ir21a*^Δ^*^1^* could also disrupt the nearby *chitin deacetylase 5* (*cda5*) gene (Supp. Fig. 2a), *Ir21a*-specific rescue experiments were performed to confirm all defects reflected the loss of *Ir21a* activity (see below).

**Figure 2:**
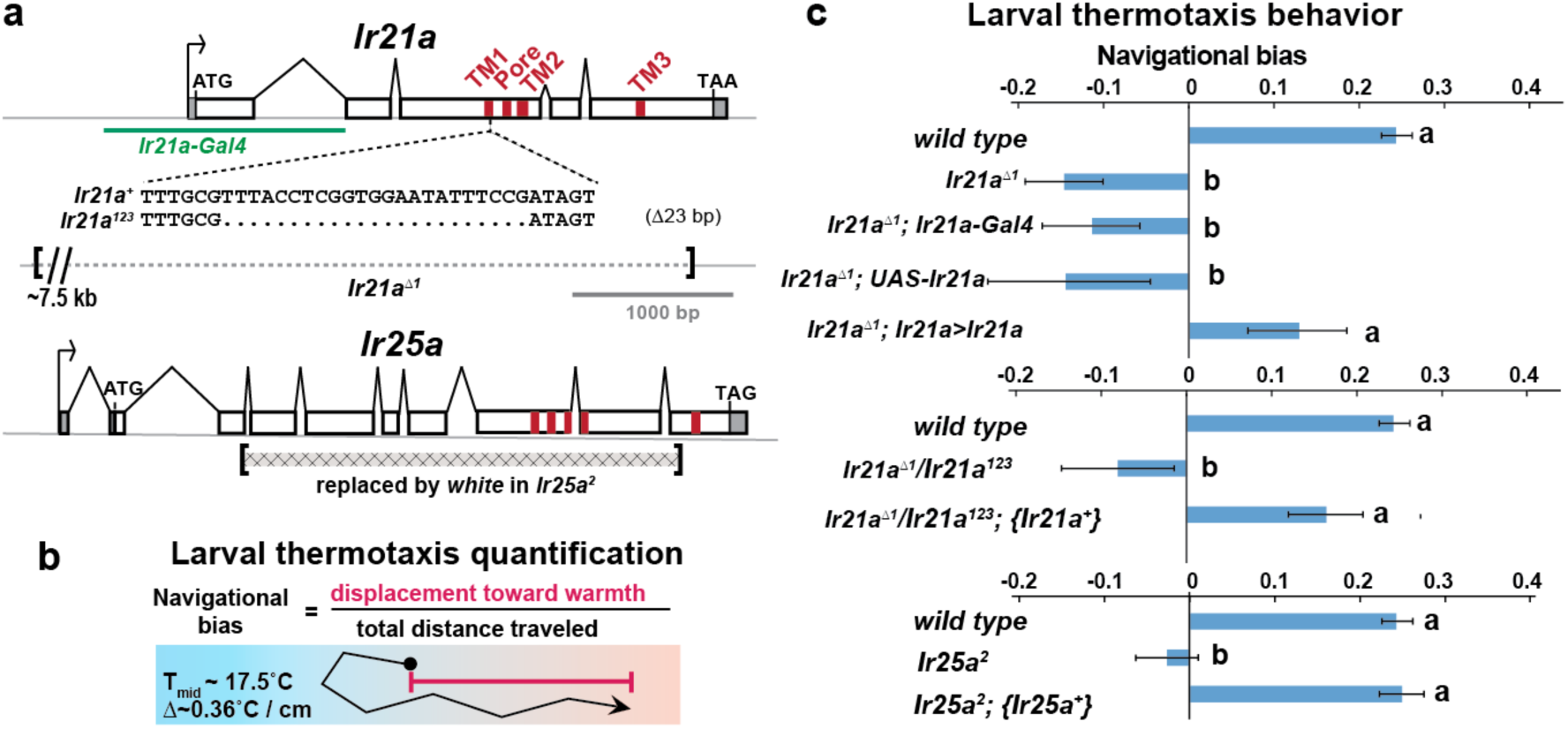
Larval cool avoidance requires *Ir21a* and *Ir25a*. a) Sequences alterations in *Ir21a* and *Ir25a* alleles. *Ir21a* regulatory sequences present in *Ir21a-Gal4* are denoted in green and regions encoding transmembrane domains (TMs) and pore region in red. b) Thermotaxis is quantified as navigational bias. Larval cool avoidance was assessed on a ~0.36°C/cm gradient extending from ~13.5°C to ~21.5°C, with a midpoint of ~17.5°C. c) Cool avoidance requires *Ir21a* and *Ir25a. Ir21a>Ir21a* denotes a wild type *Ir21a* transcript expressed under *Ir21a-Gal4* control. {*Ir21a^+^*} and {*Ir25a^+^*} denote wild type genomic rescue transgenes. Letters denote statistically distinct categories (alpha=0.05; Tukey HSD). *wild type*, n=836 animals. *Ir21a*^Δ^*^1^*, n=74. *Ir21a*^Δ^*^1^;Ir21a-Gal4*, n=48. *Ir21a*^Δ^*^1^;UAS-Ir21a*, n=10. *Ir21a*^Δ^*^1^;Ir21a>Ir21a*, n= 88. *Ir21a*^Δ^*^1^/ Ir21a^123^*, n=71; *Ir21a*^Δ^*^1^/ Ir21a^123^;* {*Ir21a+*} n=70; *Ir25a^2^*, n =100. *Ir25a^2^;* {*Ir25a+*} n= 247.

Consistent with a critical role for *Ir21a* in larval thermotaxis, the loss of *Ir21a* function strongly disrupted larval thermotaxis. When exposed to a thermal gradient of ~0.36°C/cm, ranging from ~13.5°C to ~21.5°C, *Ir21a*^Δ^*^1^* null mutants as well as *Ir21a^123^/Ir21a*^Δ^*^1^* heterozygotes were unable to navigate away from cooler temperatures and toward warmer temperatures (Fig. 2b, 2c). These defects could be rescued by expression of a wild-type *Ir21a* transcript under *Ir21a-Gal4* control and by a wild-type *Ir21a* genomic transgene (Fig. 2c). Taken together, these results are consistent with a critical role for *Ir21a* in larval thermotaxis.

### *Ir25a* mediates larval thermotaxis and is expressed in DOCCs

As IRs commonly act in conjunction with “co-receptor” IRs, we examined the possibility that larval thermotaxis involved such additional IRs. Animals homozygous for loss-of-function mutations in two previously reported IR coreceptors, *Ir8a* and *Ir76b*, exhibited robust avoidance of cool temperatures, indicating that these receptors are not essential for this behavior (Supp Fig. 2b). By contrast, *Ir25a^2^* null mutants failed to avoid cool temperatures, a defect that could be rescued by the introduction of a transgene containing a wild type copy of *Ir25a* (Fig. 2c). Thus, *Ir25a* also participates in cool avoidance. To assess IR25a expression, larvae were stained with antisera for IR25a. Robust IR25a protein expression was detected in multiple cells in the dorsal organ ganglion, including the three *Ir21a-Gal4-expressing* DOCCs (Fig. 3a). Within DOCCs, IR25a strongly labels the “dendritic bulbs”, consistent with a role in sensory transduction. Staining was absent in *Ir25a* null mutants demonstrating staining specificity (Fig. 3b). Thus *Ir25a* is required for thermotaxis and is expressed in the neurons that drive this behavior.

**Figure 3:**
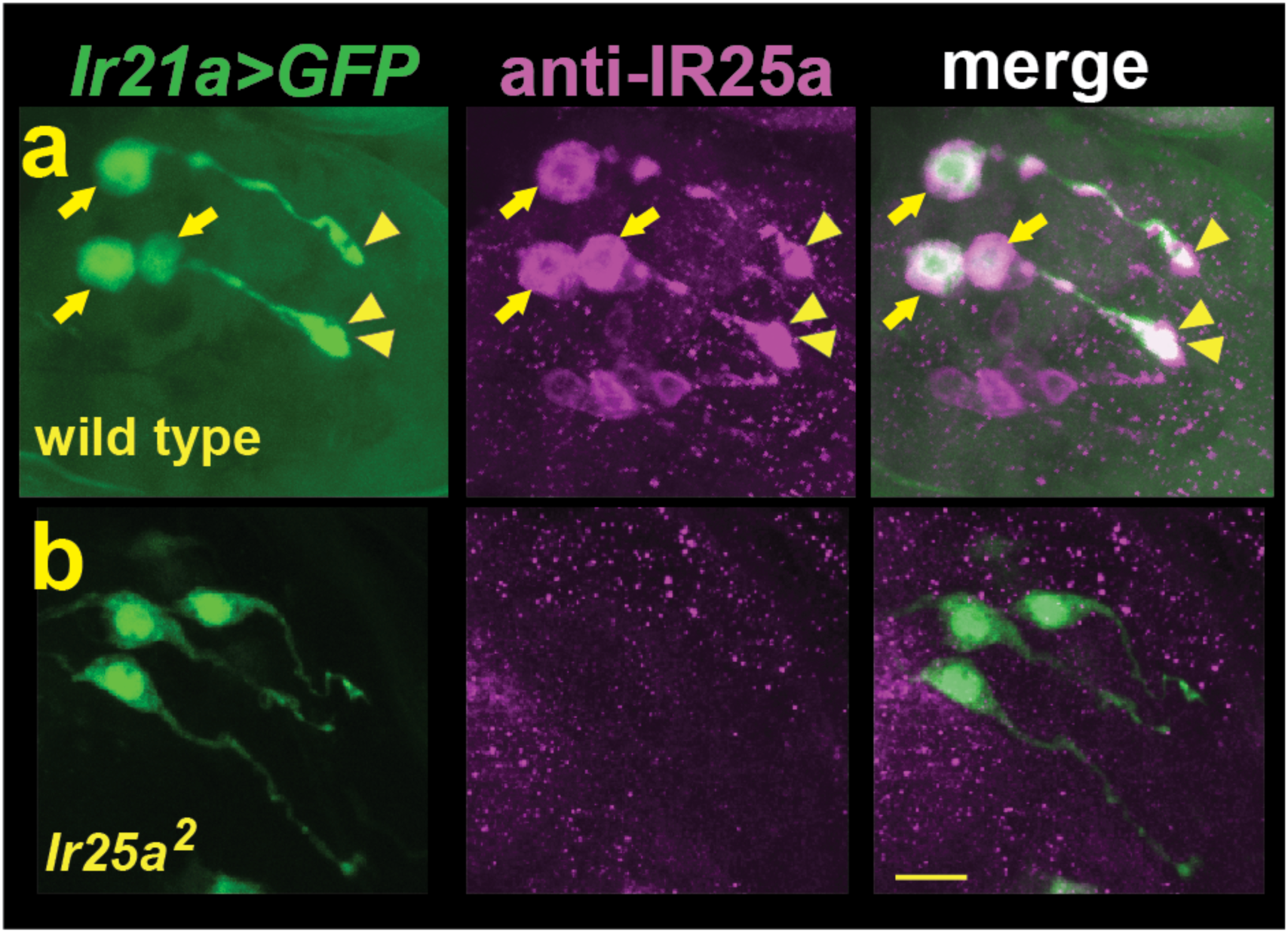
DOCCs express IR25a. a) Left panel, *Ir21a>GFP*-labeled DOCCs. Middle panel, IR25a protein expression in dorsal organ. Right panel, *Ir21a>GFP-*labeled DOCCs express IR25a protein. Arrows denote DOCC cell bodies and arrowheads DOCC dendritic bulbs. b) IR25a immunostaining is not detected in *Ir25a^2^* null mutants. Scale bar, 10 microns.

### *Ir21a* and *Ir25a* are required for cool detection by DOCCs

To assess whether *Ir21a* and *Ir25a* contribute to cool detection by the DOCCs, DOCC cool-responsiveness was examined using the genetically encoded calcium sensor GCaMP6m. Consistent with a role for *Ir21a* in cool responses, DOCCs exhibited strongly reduced responses to cooling in *Ir21a*^Δ^*^1^* deletion mutants, and this defect was robustly rescued by expression of an *Ir21a* transcript in the DOCCs using *R11F02-Gal4* (Fig. 4a-e, 4h). Similarly, DOCC thermosensory responses were greatly reduced in *Ir25a* mutants, a defect that was rescued by a wild type *Ir25a* transgene (Fig. 4f-h). Together these data demonstrate a critical role for *Ir21a* and *Ir25a* in the detection of cooling by the DOCCs.

**Figure 4:**
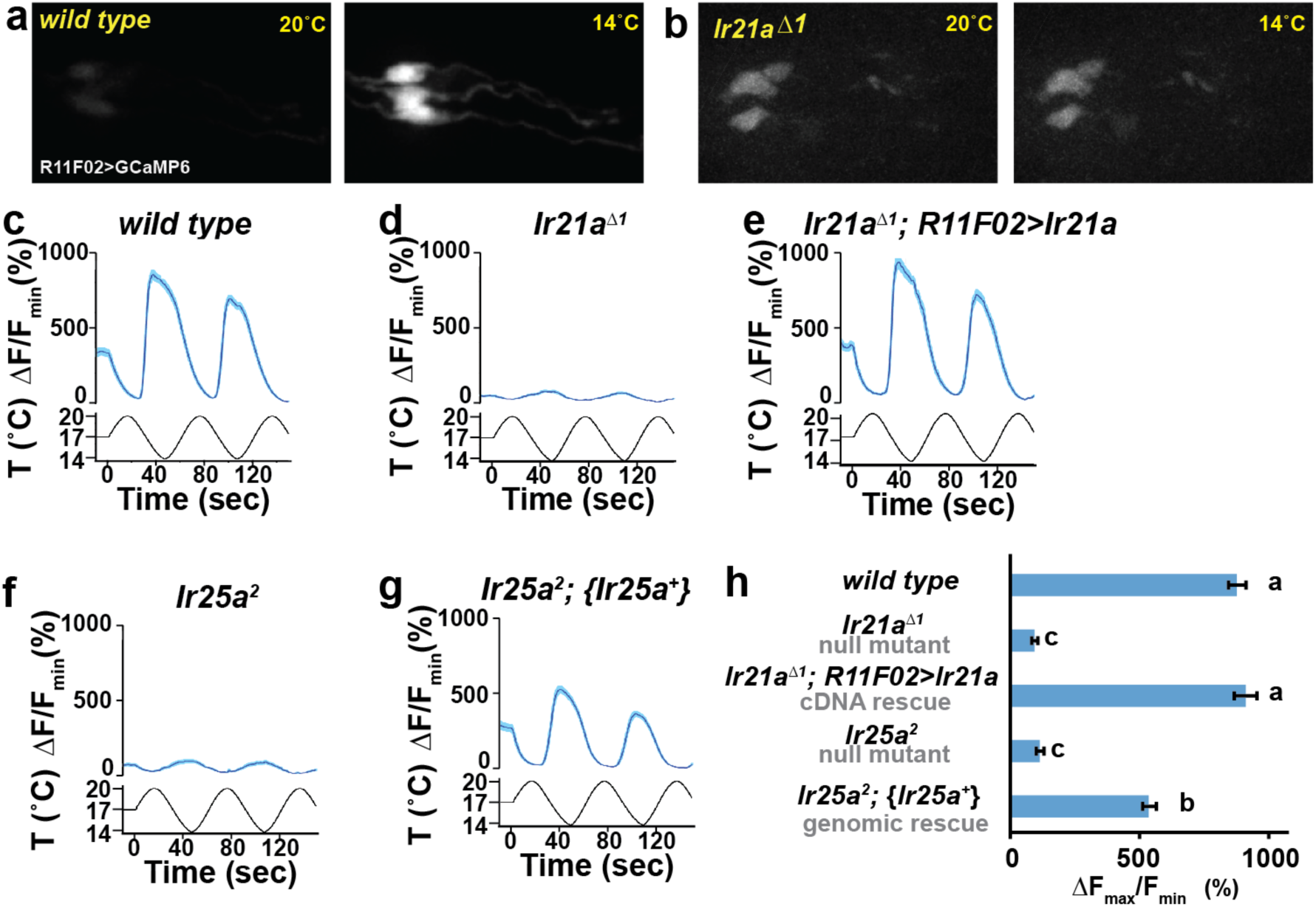
DOCC cool responses require *Ir21a* and *Ir25a*. DOCC responses monitored using *R11Fo2>GCaMP6m*. DOCCs exhibit robust cool-responsive increases in fluorescence (a,c), which are dramatically reduced in *Ir21a* (b,d) and *Ir25a* (f) mutants. e) *Ir21a* transcript expression under *R11F02-Gal4* control rescues the *Ir21a* mutant defect. g) Introduction of an *Ir25a* genomic rescue transgene rescues the *Ir25a* mtuant defect. h) Ratio of fluorescence at 14°C versus 20°C. Letters denote statistically distinct categories, alpha=0.01, Tukey HSD. Scale bars, 10 microns. Traces, average +/− SEM. *wild type*, n=33 cells. *Ir21a*^Δ^*^1^*, n= 58. *Ir21a*^Δ^*^1^; R11Fo2>Ir21a*, n=32. *Iv25a^2^*, n=43. *Ir25a^2^;* {*Ir25a^+^*}, n=30.

Prior work has suggested that three TRP channels, Brivido-1, Brivido-2 and Brivido-3, work together to mediate cool sensing in adult thermosensors (Gallio et al., 2011). Putative null mutations are available for two of these genes, *brv1* and *brv2*, and we used these alleles to test the potential role of Brivido function in DOCC cool sensing (Gallio et al., 2011). Although *brv1* mutant showed defects in thermotactic behavior, DOCC responses to cooling appeared unaffected in *brv1* mutants (Supp. Fig. 4a, 4b). *brv2* nulls exhibited no detectable thermotaxis defects (Supp. Fig. 4a). Thus we detect no role for these receptors in cool sensing by the DOCCs.

### Ectopic IR21a expression confers cool-sensitivity in an *Ir25a-*dependent fashion

The requirement for *Ir21a* and *Ir25a* in DOCC-mediated cool sensing raised the question of whether ectopic expression of these receptors could confer cool-responsiveness upon a cell, as might be predicted for a cool receptor. Attempts to express IR21a and IR25a together or separately in heterologous cells, including S2 cells, *Xenopus* oocytes and HEK cells, failed to yield detectable cool-activated currents, as did attempts to ectopically express them separately and together in *Drosophila* chemosensory neurons (G.B., L.N. and P.G, unpublished). However, ectopic expression of IR21a in one set of neurons in the adult, Hot Cell thermoreceptors in the arista that normally respond to warming rather than cooling, did confer cool-sensitivity.

The adult arista contains three warmth-activated thermosensory neurons, termed Hot Cells (or HC neurons) (Gallio et al., 2011). We found that forced expression of IR21a in the HC neurons could significantly alter their response to temperature. As previously reported (Gallio et al., 2011), wild-type HC neurons respond to warming with robust increases in intracellular calcium and to cooling with decreases in intracellular calcium, as reflected in temperature-dependent changes in GCaMP6m fluorescence (Fig. 5a, 5c). In contrast, HC neurons in which IR21a is expressed under the control of a pan-neuronal promoter (*N-syb>Ir21a* animals) frequently exhibited elevations in calcium not only in response to warming, but also at the coolest temperatures (Fig. 5b, 5d, 5f). Thus, ectopic IR21a expression causes HC neurons to respond to cooling as well as warming.

**Figure 5:**
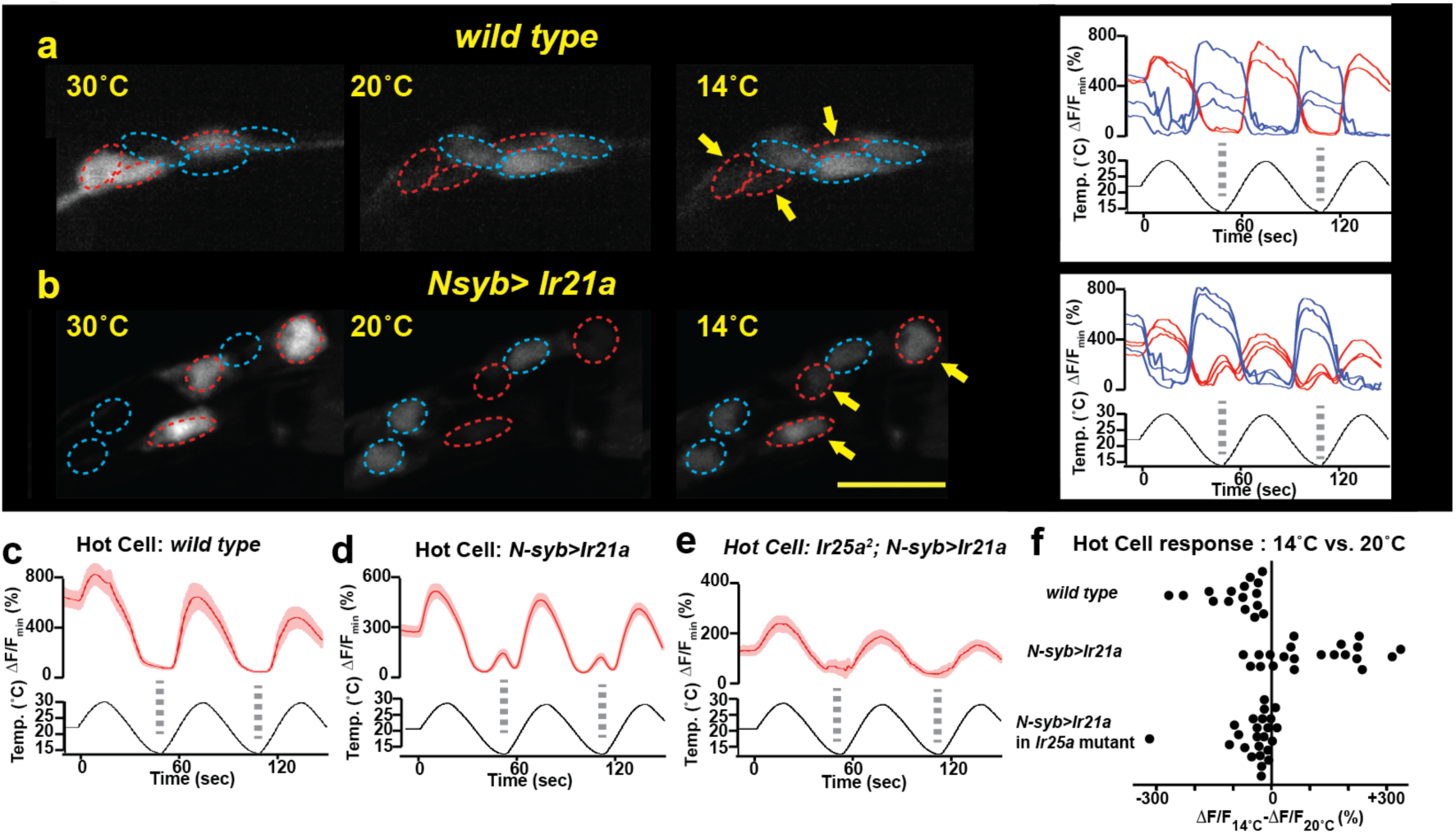
IR21a expression confers cool-sensitivity upon warmth-responsive Hot Cell neurons. a,b) Temperature responses of *wild type* (a) or *N-syb>Ir21a*-expressing (b) thermoreceptors in the adult arista, monitored with *N-syb>GCaMP6m*. Cell bodies of warmth-responsive Hot Cells outlined in red and cool-responsive Cold Cells in blue. Arrows highlight Hot Cells at 14°C. Traces of individual Hot Cell and Cold Cell responses shown at right. c-e) Fluorescence of Hot Cells in response to sinusoidal 14°C to 30°C temperature stimulus, quantified as percent ΔF/F_min_. Dotted lines denote temperature minima. Traces, average +/− SEM. f) Difference between ΔF/F_min_ at 14°C and ΔF/F_min_ at 20°C for each cell imaged. Responses of *N-syb>Ir21a* cells were statistically distinct from both *wild type* and *Ir25a^2^;N-syb>Ir21a* (p<0.01, Steel-Dwass test). Scale bar, 10 microns. *wild type*, n= 16 cells. *N-syb>Ir21a*, n= 16. *Ir25a^2^; N-syb>Ir21a*, n= 20.

As *Ir21a*-dependent cool detection in the DOCCs relies upon *Ir25a*, we examined the requirement for *Ir25a* in IR21a-mediated cool activation of the HC neurons. Consistent with previously reported IR25a expression in the arista (Benton et al., 2009), we observed robust IR25a protein expression in the HC neurons (Supp. Fig. 5a, 5b). Consistent with a role for *Ir25a* in *Ir21a*-mediated cool-responsiveness, ectopic IR21a expression failed to drive significant HC neuron cool responses in *Ir25a* mutants (Fig. 5e, 5f). Thus, IR21a can confer cool-sensitivity upon an otherwise warmth-responsive neuron in an *Ir25a*-dependent fashion. Similar cool sensitivity is observed when IR21a is ectopically expressed under the control of an HC-specific promoter (*HC>Ir21a*, Supp. Fig. 5c, 5d). Together, these data demonstrate that IR21a expression can confer cool-sensitivity on the normally warmth-sensitive HC neurons in an *Ir25a*-dependent fashion.

## DISCUSSION

These data demonstrate that the Ionotropic Receptors IR21a and IR25a have critical roles in thermosensation in *Drosophila*, mediating cool detection by the larval dorsal organ cool cells (DOCCs) and the avoidance of cool temperatures. Combinations of IRs have been previously found to contribute to a wide range of chemosensory responses, including the detection of acids and amines (Rytz et al., 2013). These findings extend the range of sensory stimuli mediated by these receptor combinations to cool temperatures.

The precise nature of the molecular complexes that IRs form is not well understood. IR25a has been shown to act with other IRs in the formation of chemoreceptors, potentially as hetero-multimers (Rytz et al., 2013). This precedent raises the appealing possibility that IR25a might form heteromeric thermoreceptors in combination with IR21a. However, the inability to readily reconstitute cool-responsive receptor complexes in heterologous cells suggests that the mechanism by which these receptors contribute to cool responsiveness is likely to involve additional molecular co-factors. It is interesting to note that the range of cell types in which ectopic IR21a expression confers cool-sensitivity is so far restricted to neurons that are already respond to temperature. This observation suggests the existence of additional co-factors or structures in these thermosensory cells that are critical for IR21a and IR25a to function. Recently, IR25a was implicated in resetting of the circadian clock by increases in external temperature (Chen et al., 2015). However, misexpression of IR25a in heterologous neurons on its own conferred only very low sensitivity responses to temperature changes (Chen et al., 2015), raising the intriguing possibility that – analogous to cool-sensing – IR25a acts with another IR to mediate detection of temperature increases.

While the present study focuses on the role of IR21a and IR25a in larval thermosensation, it is interesting to note that expression of both IR21a and IR25a has been detected in the thermoreceptors of the adult arista (Benton et al., 2009). Thus related mechanisms could contribute to thermosensory responses not only in the DOCCs, but also in other cellular contexts and life stages. Moreover, the presence of orthologs of IR21a and IR25a across a range of insects (Croset et al., 2010) raises the possibility that these IRs, along other members of the IR family, constitute a family of deeply-conserved thermosensors.

### Material and Methods

**Fly strains**. *Ir25a^2^* (Benton et al., 2009), *BAC*{*Ir25a^+^*} (Benton et al., 2009), *Ir8a^1^* (Benton et al., 2009), *Ir76b^1^* (Zhang et al., 2013), *Ir76b^2^* (Zhang et al., 2013), *R11F02-Gal4* (Klein et al., 2015), *brv1^L653stop^* (Gallio et al., 2011), *brv2^w205stop^* (Gallio et al., 2011), *HC-Gal4* (Gallio et al., 2011), *UAS-GCaMP6m* (P{20XUAS-IVS-GCaMP6m}attp2 and P{20XUAS-IVS-GCaMP6m}attp2attP40 (Chen et al., 2013)), *UAS-GFP* (p{10X UAS-IVS-Syn21-GFP-p10}attP2 (Pfeiffer et al., 2012)), *nSyb-Gal4* (P{GMR57c10-Gal4}attP2, (Pfeiffer et al., 2012)), and y1 P(act5c-cas9, w+) M(3xP3-RFP.attP)ZH-2A w* (Port et al., 2014) were previously described.

In *Ir21a-Gal4*, sequences from-606 to +978 with respect to the *Ir21a* translational start site (chromosome 2L: 24173 – 25757, reverse complement) lie upstream of Gal4 protein-coding sequences. *UAS-Ir21a* contains the *Ir21a* primary transcript including introns (chromosome 2L: 21823–25155, reverse complement) placed under UAS control. The {*Ir21a+*} genomic rescue construct contains sequences from −1002 to +4439 with respect to the *Ir21a* translational start site (chromosome 2L: 26153–20712).

*Ir21a*^Δ^*^1^* was generated by FLP-mediated recombination between two FRT-containing transposon insertions (PBac{PB}c02720 and PBac{PB}c04017) as described (Parks et al., 2004). *Ir21a^123^* was generated by transgene-based CRISPR-mediated genome engineering as described (Port et al., 2014), with an Ir21a-targeting gRNA (5’-CTGATTTGCGTTTACCTCGG) expressed under U6–3 promoter control (dU6–3:gRNA) in the presence of *act-cas9* (Port et al., 2014).

**Behaviour**. Thermotaxis of early 2^nd^ instar larvae was assessed over a 15 min period on a temperature gradient extending from 13.5 to 21.5°C over 22 cm (~0.36 °C/cm) as described (Klein et al., 2015).

**Calcium imaging**. Calcium imaging was performed as previously described for larvae (Klein et al., 2015). Pseudocolor images were created using the 16_colors lookup table in ImageJ 1.43r. Adult calcium imaging was performed as described for larvae (Klein et al., 2015), with modifications to the temperature stimulus and sample preparation approach. Adult temperature stimulus ranged from 14°C to 30°C. Intact adult antennae with aristae attached were dissected and placed in fly saline (110 mM NaCl, 5.4 mM KCl, 1.9 mM CaCl2, 20 mM NaHCO_3_, 15 mM tris(hydroxymethyl)aminomethane (Tris), 13.9 mM glucose, 73.7 mM sucrose, and 23 mM fructose, pH 7.2, (Brotz and Borst, 1996)) on a large cover slip (24 mm × 50 mm) and then covered by a small cover slip (18 mm × 18 mm). The large cover slip was placed on top of a drop of glycerol on the temperature control stage.

**Immunohistochemistry**. Immunostaining was performed as described (Kang et al., 2012) using rabbit anti-Ir25a (1:100; (Benton et al., 2009)), mouse anti-GFP (1:200; Roche), goat anti-rat Cy3 (1:100; Jackson ImmunoResearch), donkey anti-mouse FITC (1:100; Jackson ImmunoResearch).

**Author contributions:** L.N., M.K., G.B., R.B., A.D.T. and P.A.G. designed experiments. L.N. performed molecular genetics, behavior, immunohistochemistry and calcium imaging, M.K. performed behavior and calcium imaging, K.S. and E.C.C. performed molecular genetics, G.B. performed physiology, L.N., M.K., R.B., A.D.T.S., and P.A.G. wrote the paper.

## Acknowledgements

We thank Rachelle Gaudet and Linda Huang for comments on the manuscript, Peter Bronk for advice on physiology, Adam Kaplan for creating *Ir21a-Gal4*, and the Bloomington Stock Center for fly strains. Supported by a grant from the National Institute of Neurological Disorders and Stroke (F32 NS077835) to M.S., European Research Council Starting Independent Researcher and Consolidator Grants (205202 and 615094) to R.B., and the National Institute of General Medical Sciences (P01 GM103770) to A.D.T.S. and P.A.G.**Competing interests**: The authors have no competing interests.

---

**Supplementary Figure 1:**
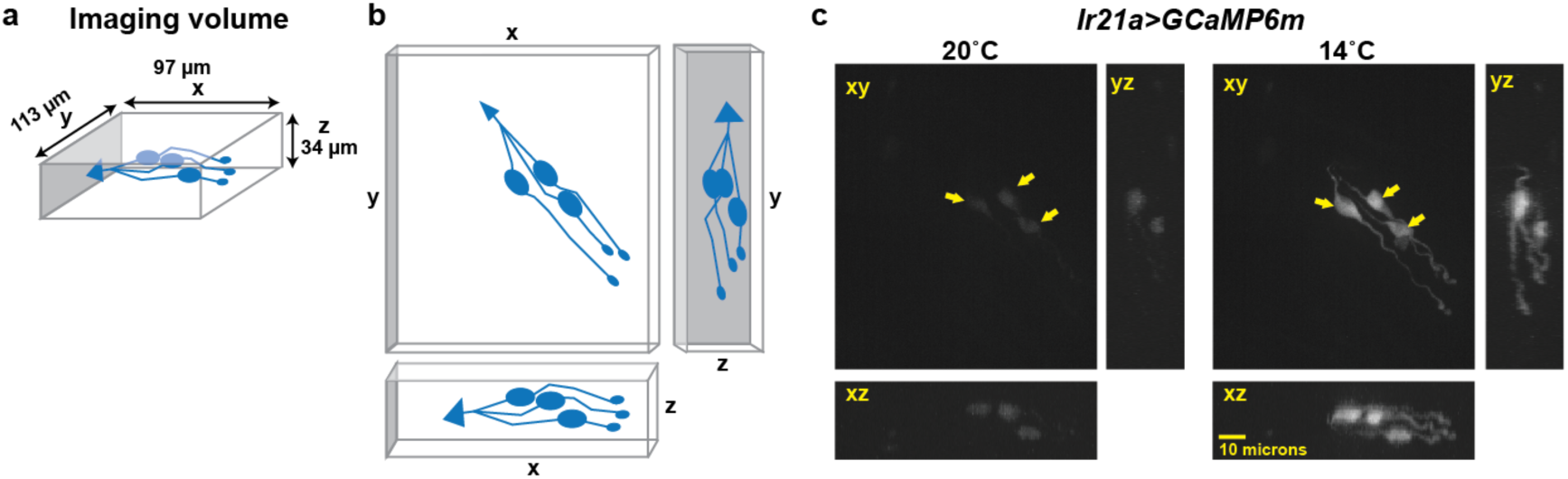
Calcium-imaging data are obtained as a three-dimensional imaging stack. a) Dimensions of imaging volume. DOCCs depicted in blue. b, Maximum intensity projections used for visualizing fluorescence intensity. c, Representative image of maximum intensity projections of *Ir21a>GCaMP6m*-labeled DOCCs. DOCC cell bodies remain within imaging field throughout.

**Supplementary Figure 2:**
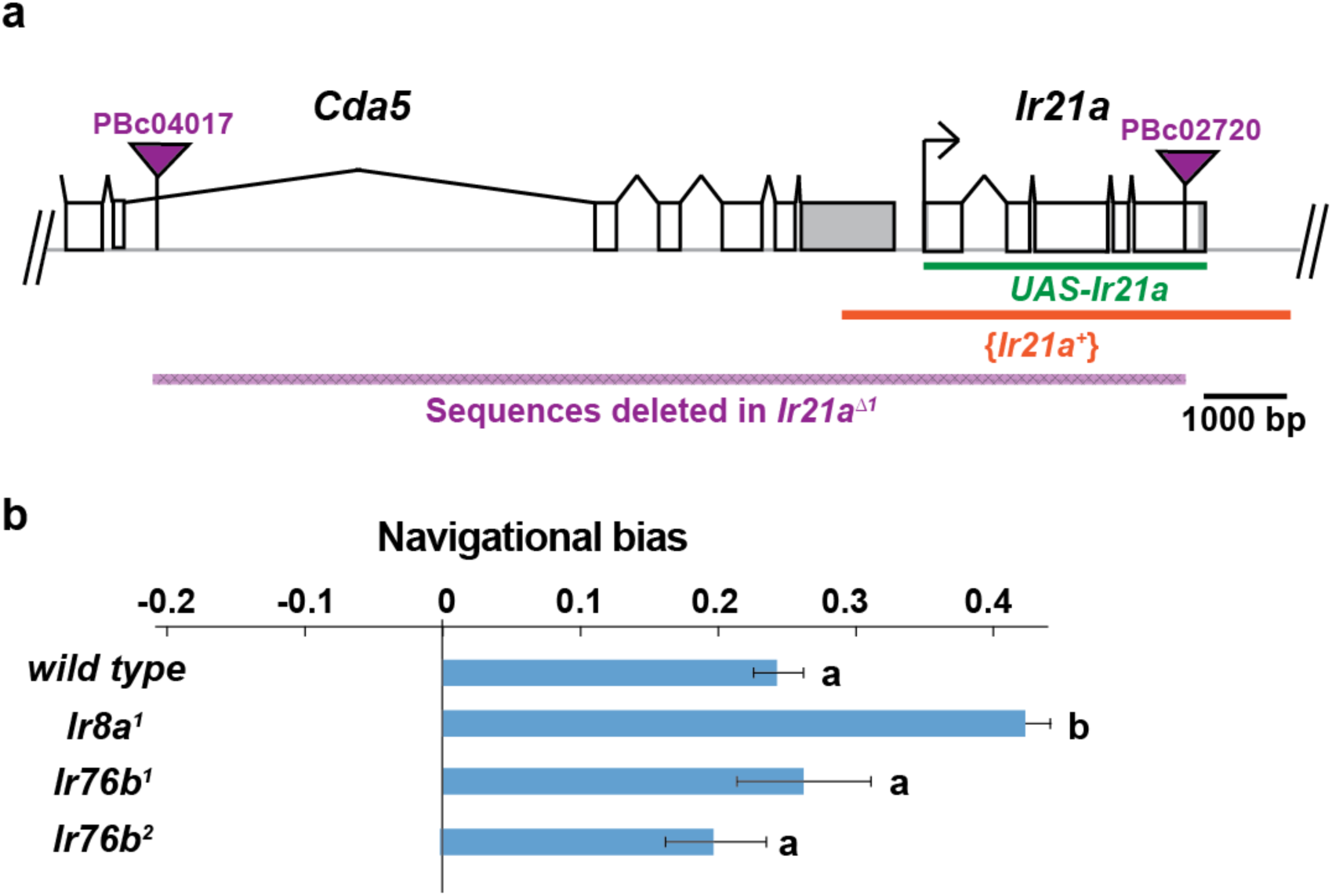
Structure of *Ir21a* locus and analysis of thermotaxis in *Ir8a* and *Iry6b* mutants. a) *Cda5/Ir21a* genomic region, denoting positions of the FRT-containing transposon insertions used to generate *Ir21a*^Δ^*^1^* (PBc04017 and PBc02720), the sequences deleted in *Ir21a*^Δ^*^1^*, the *Ir21a* sequences present in the *UAS-Ir21a* rescue construct and the sequences present in the {*Ir21^+^*} genomic rescue construct. Untranslated regions are in gray. b) Larval thermotaxis of *Ir8a* and *Iry6b* mutants quantified as navigational bias. Neither *Ir8a* nor *Iry6b* is required for cool avoidance; *Ir8a* mutants show enhanced cool avoidance compared to *wild type*. Letters denote statistically distinct categories (alpha=0.05; Tukey HSD). *wild type*, n=836 animals. *Ir8a*, n=166; *Ir76b^1^*, n=96, *Ir76b^2^*, n= 100.

**Supplementary Figure 4:**
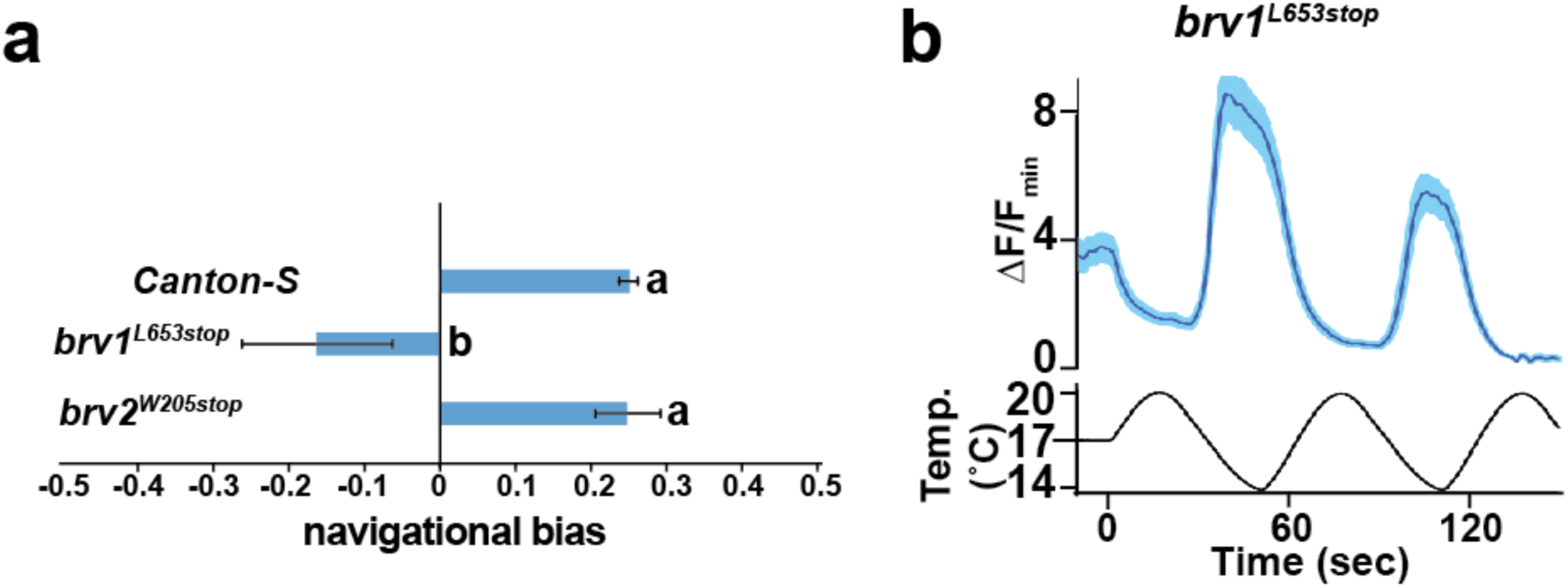
Analysis of putative null mutants of *brv1* and *brv2*. a) *brv1* but not *brv2* mutants exhibit defects in larval cool avoidance. Thermotaxis quantified as navigational bias. Letters denote statistically distinct categories (alpha=0.05; Tukey HSD). *wild type*, n=836 animals. *brv1^L653St^°^p^*, n =43. *brv2^W205St^°^p^*, n =99. b) *Ir21a>GCaMP6m*-labelled DOCCs respond to cooling in *brv1^L653St^°^p^* mutants. n= 35 cells.

**Supplementary Figure 5:**
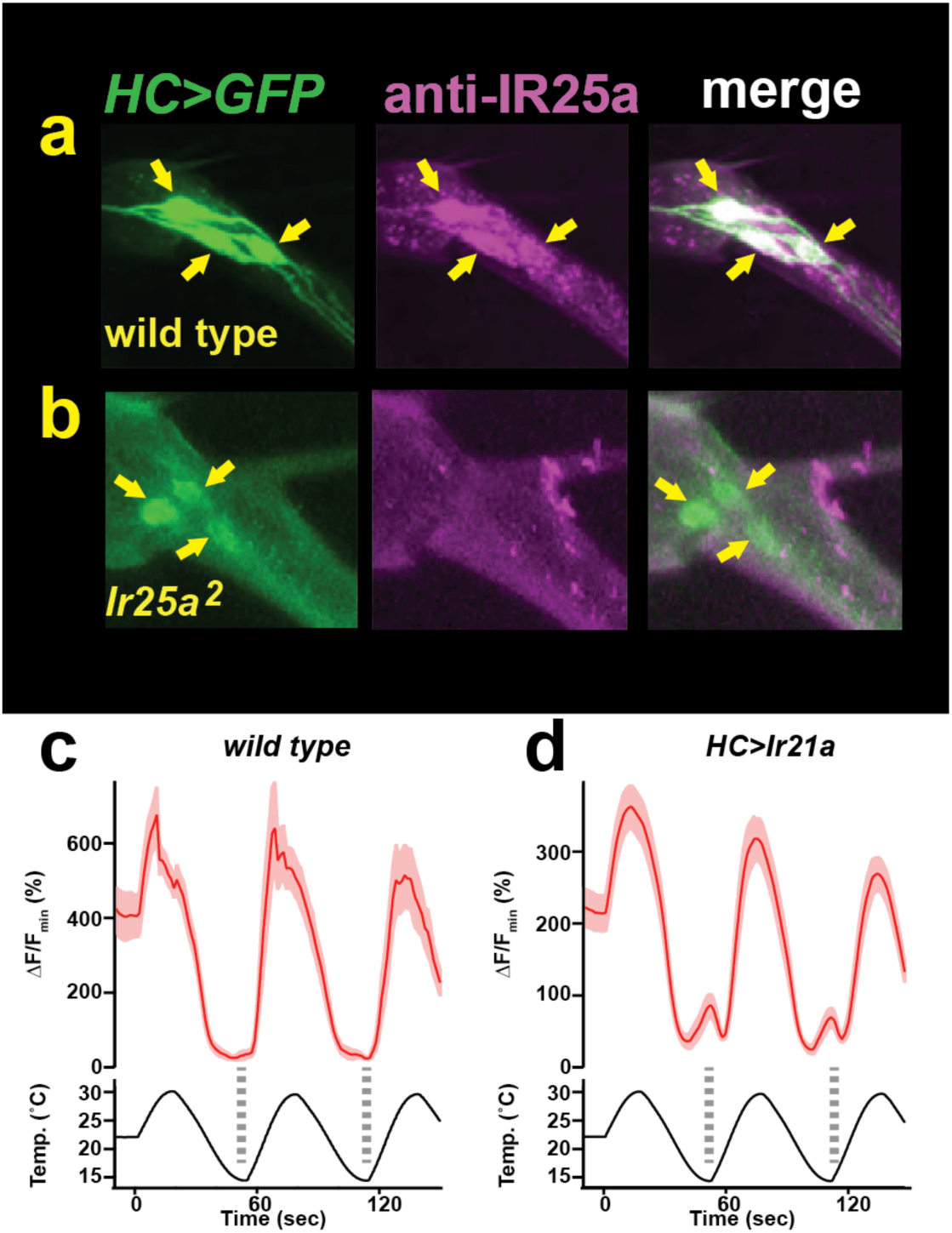
Hot Cell neurons express Ir25a protein. a,b) Left panel, HC>GFP-labeled Hot Cell neurons. Middle panel, IR25a immunostaining. Right panel, HC>GFP and Ir25a are co-expressed in the Hot Cell neurons. Arrows indicate Hot Cell neuron cell bodies. Specific IR25a immunostaining is observed in wild type HC neurons, but is absent in *Ir25a* null mutants (b). c, d) Temperature responses of *wild type* (c) or *HC>Ir21a-*expressing (d) thermoreceptors in the adult arista, monitored using *HC>GCaMP6m*. Dotted lines denote temperature minima. Traces, average +/− SEM. n=4 wild type cells, n=25 cells *HC>IR21a*.

